# Effect of the SARS-CoV-2 Delta-associated G15U mutation on the s2m element dimerization and its interactions with miR-1307-3p

**DOI:** 10.1101/2023.02.10.528014

**Authors:** Caylee L. Cunningham, Caleb J. Frye, Joseph A. Makowski, Adam H. Kensinger, Morgan Shine, Ella J. Milback, Patrick E. Lackey, Jeffrey D. Evanseck, Mihaela-Rita Mihailescu

## Abstract

The stem loop 2 motif (s2m), a highly conserved 41-nucleotide hairpin structure in the severe acute respiratory syndrome coronavirus 2 (SARS-CoV-2) genome, serves as an attractive therapeutic target that may have important roles in the virus life cycle or interactions with the host. However, the conserved s2m in Delta SARS-CoV-2, a previously dominant variant characterized by high infectivity and disease severity, has received relatively less attention than that of the original SARS-CoV-2 virus. The focus of this work is to identify and define the s2m changes between Delta and SARS-CoV-2 and subsequent impact of those changes upon the s2m dimerization and interactions with the host microRNA miR-1307-3p. Bioinformatics analysis of the GISAID database targeting the s2m element reveals a greater than 99% correlation of a single nucleotide mutation at the 15^th^ position (G15U) in Delta SARS-CoV-2. Based on ^1^H NMR assignments comparing the imino proton resonance region of s2m and the G15U at 19°C, we find that the U15-A29 base pair closes resulting in a stabilization of the upper stem without overall secondary structure deviation. Increased stability of the upper stem did not affect the chaperone activity of the viral N protein, as it was still able to convert the kissing dimers formed by s2m G15U into a stable duplex conformation, consistent with the s2m reference. However, we find that the s2m G15U mutation drastically reduces the binding affinity of the host miR-1307-3p. These findings demonstrate that the observed G15U mutation alters the secondary structure of s2m with subsequent impact on viral binding of host miR-1307-3p, with potential consequences on the immune response.

## Introduction

The effects of the coronavirus disease 2019 (COVID-19) pandemic continue to impact the world, as the virus responsible for the outbreak, SARS-CoV-2, continues to acquire advantageous mutations, resulting in new viral variants. To date, SARS-CoV-2 has resulted in over 668 million cases and 6.73 million deaths worldwide, underscoring the need to characterize the viral life cycle and interactions with the host so that future pandemics may be thwarted (CDC 2020b). Although some variants of SARS-CoV-2 were short lived, several created new waves of infections. These variants exhibited heightened levels of “immune escape” as well as changes in symptom severity (CDC 2020b, 2020a). One of these variants, identified as the Delta variant, originated in India with the earliest documented case in October of 2020. Delta was subsequently declared a variant of concern (VOC) by the World Health Organization in May 2021 and became the dominant variant of the virus soon after. In November of 2021, Delta cases began decreasing as a new prominent variant, Omicron, emerged (CDC 2020b; Yaniv et al. 2022). Omicron was quickly declared a VOC on November 26, 2021 and has since mutated into several sublineages, with recent work suggesting that Omicron gained several mutations throughout its genome, aiding in transmissibility (CDC 2020a; Jung et al.; Meng et al. 2022). As new major SARS-CoV-2 variants and sublineages emerged, their transmissibility increased while the disease severity fluctuated, suggesting the virus is adapting to increase its viral spread and longevity (CDC 2022). Although Omicron is presently the most prevalent variant, it was not able to fully out-compete its VOC predecessor, as Delta cases collected as late as December 2022 and January 2023 are still reported in the various parts of the world (e.g., EPI_ISL_16252097 in the USA; EPI_ISL_16279089 and EPI_ISL_16279092 in Botswana; EPI_ISL_16823192 and EPI_ISL_16823199 in India). This is unlike the eradication of the previous Gamma and Beta variants, making the Delta variant even more interesting to study due to its viability (Yaniv et al. 2022).

SARS-CoV-2 variants are classified by advantageous mutations to the spike (S) protein, particularly those which increase binding affinity to the angiotensin converting enzyme 2 (ACE2) receptor, the viral entry point into the host cell (Yan et al. 2020). However, in addition to open reading frames, the SARS-CoV-2 genome contains both 5’- and 3’-untranslated regions (UTRs) that harbor highly conserved structures that play vital roles in the replication of both the genomic and subgenomic RNAs (Malone et al. 2022; Lulla et al. 2021). In addition to differences in the S protein, mutations exist within the Delta variant’s 5’- and 3’-UTRs. We focused on the 3’-UTR of SARS-CoV-2 as it contains a 41 nucleotide (nt) highly conserved secondary structure termed the stem-loop II element (s2m) (Figure 1A, bottom) (Tengs et al. 2013). Previous research has demonstrated that in addition to coronaviruses, this motif is present in three other single-stranded viral families, *Astroviridae*, *Caliciviridae*, and *Picornaviridae* (Tengs and Jonassen 2016; Tengs et al. 2013). Although the exact roles of this element have not been elucidated, the high level of s2m sequence conservation within the hyper variable region of the 3’-UTR led to the belief that it is beneficial to the virus. Some of the suggested functions of the s2m include host microRNA (miR) hijacking, host protein hijacking, and involvement in viral recombination events (Robertson et al. 2005; Yeh and Contreras 2020). We previously reported that, through the presence of a palindromic sequence at the terminal loop, a kissing dimer structure forms between two s2m elements which are converted to an extended duplex structure by the viral N protein (Figure 2) (Imperatore et al. 2022). From this observation, we proposed that this process could be important in recombination and template switching, leading to the adaptation of new mutations to come (Muriaux et al. 1996; Shetty et al. 2010; Imperatore et al. 2022). Additionally, we have shown that the s2m is able to bind up to two copies of the host miR-1307-3p (Imperatore et al. 2022). The binding of miRs to the virus can be beneficial to enhance viral replication; however, this interaction may also be a part of the host’s immune response to the viral infection (Arisan et al. 2020; Balmeh et al. 2020).

**Figure 1.**
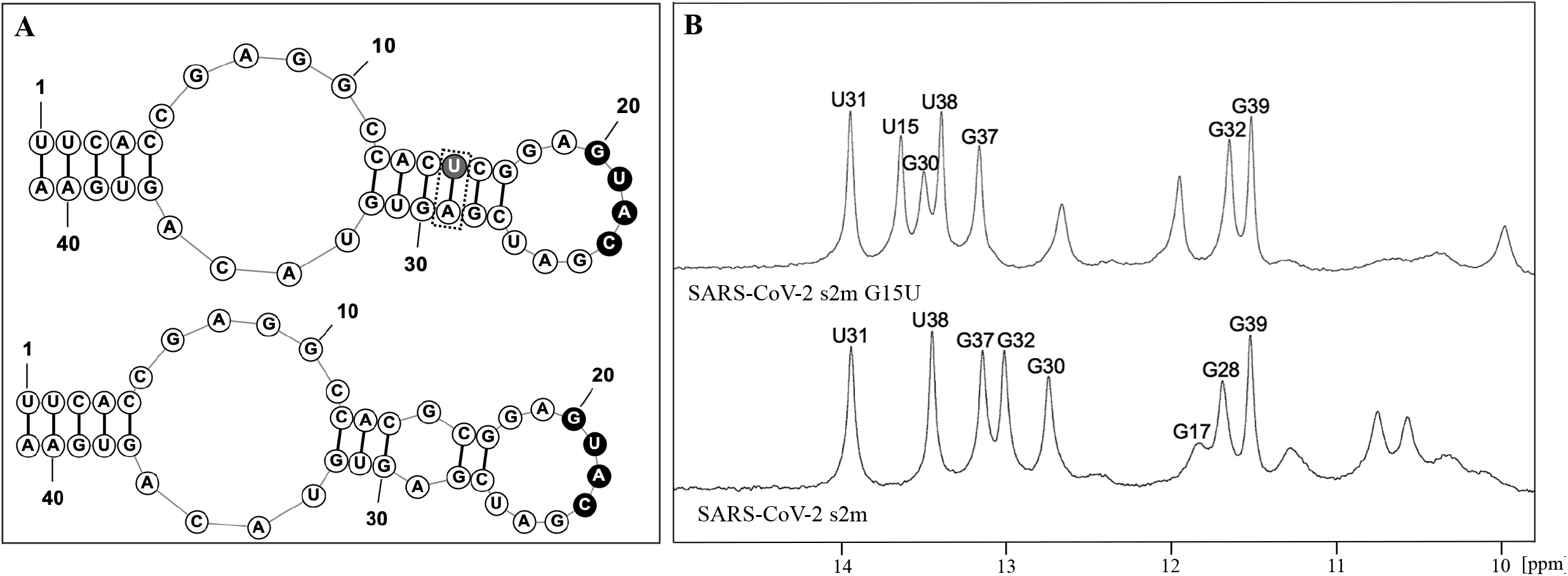
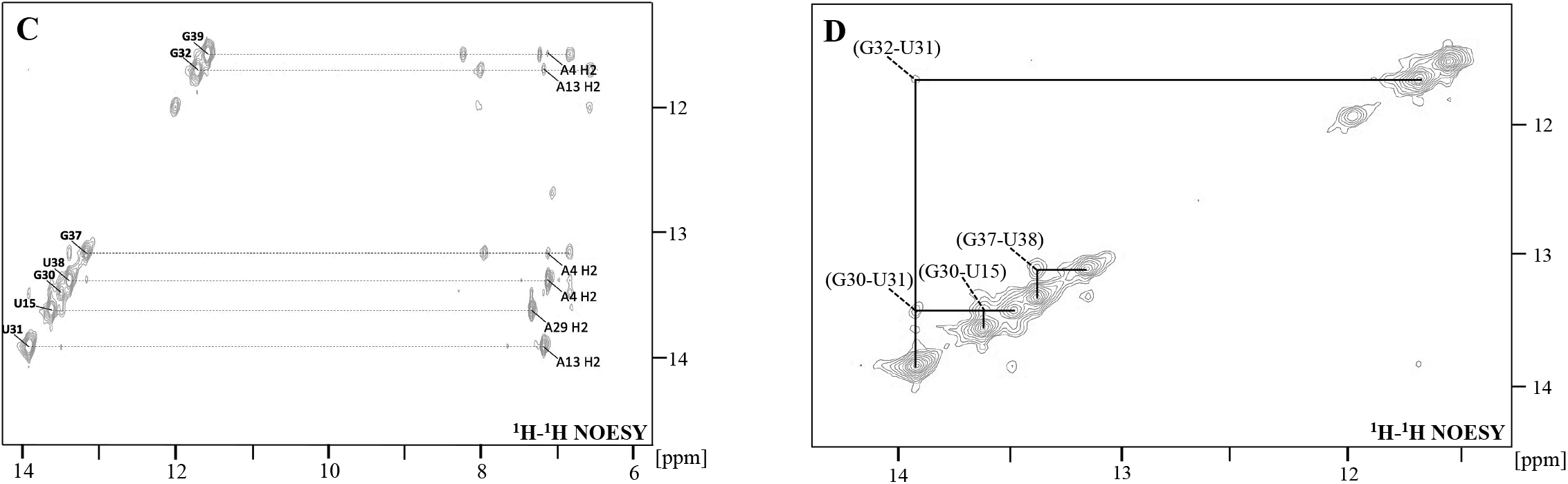
**A.** RNAStructure representation of the secondary structures of the SARS-CoV-2 reference s2m (bottom) and G15U mutation (top). The closing of the U15 and A29 base pair is indicated in the top s2m G15U by the dashed box. **B.** One-dimensional H NMR spectroscopy of the SARS-CoV-2 s2m G15U (top) and SARS-CoV-2 reference s2m (bottom). The labeled resonances were assigned previously for the reference s2m and in the current work for s2m G15U (Imperatore et al. 2022; Wacker et al. 2020). **C.** ^1^H-^1^H NOESY spectra of the imino proton resonances of s2m G15U with labeled NOEs between base paired U imino and A H2 and between G imino protons and neighboring A H2. **D.** ^1^H-^1^H NOESY spectra of the imino proton resonances for the s2m G15U with labeled NOE between neighboring intra-strand and inter-strand Gs and Us.

**Figure 2.**
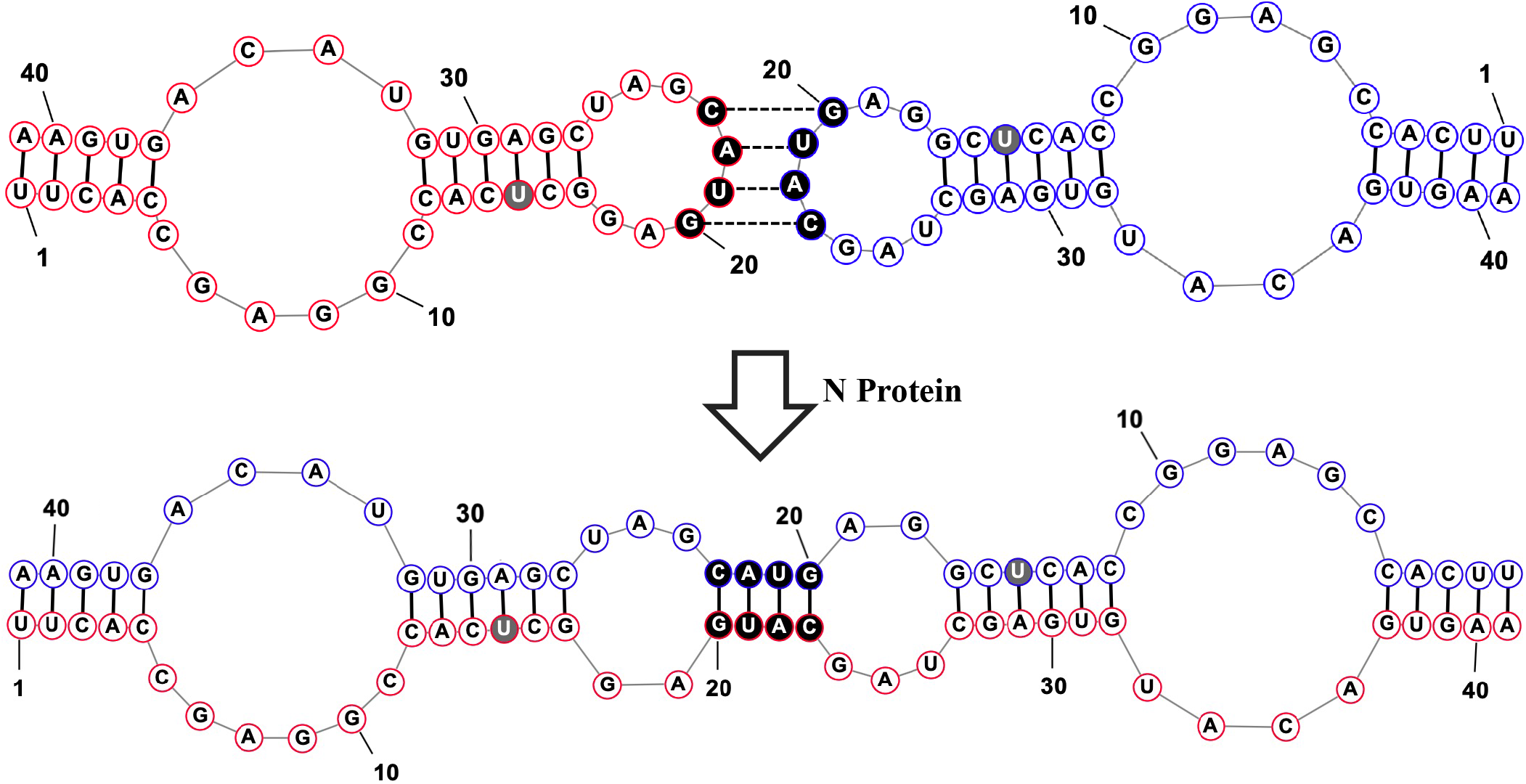
RNAStructure depiction of two s2m G15U elements (shown in red and blue for clarity) forming a kissing dimer structure through their palindromic sequences, GUAC (colored circles). The N protein then acts as a chaperone to convert the kissing dimer intermediate to an extended duplex, a more stable structure. The G15U mutation in the upper stem is highlighted by the filled circle.

Notably, only a two-nucleotide difference exists between the s2m of SARS-CoV-2 and SARS-CoV, the related virus responsible for the 2002-2003 outbreak of “SARS.” One of those mutations, G>U at position 31 (G31U) of the s2m (29,758 in the reference SARS-CoV-2 genome, EPI_ISL_402123), has previously been conserved as a G in all other reported s2m elements, and our earlier study revealed that G31U is the causative mutation between differences in dimerization properties in SARS-CoV-2 s2m and SARS-CoV s2m, as well as in miR-1307-3p binding (Imperatore et al. 2022; Zhao et al. 2020). Moreover, our molecular dynamics (MD) simulation study showed that these dimerization differences are likely caused by increases in structural flexibility of the terminal loop in SARS-CoV-2 s2m compared to the rigid GNRA-like pentaloop motif found in the SARS-CoV s2m (Kensinger et al. 2022). In SARS-CoV-2 s2m, an expansion to a “nonaloop” was computed to confer an entropic penalty, potentially explaining the reduction in kissing complex formation, the intermediate step in duplex formation (Kensinger et al. 2022).

Knowing the drastic impact of a single nucleotide mutation on the structure of SARS-CoV-2 s2m, we conducted a bioinformatics analysis ranging from April 2020 to October 2021 to identify new mutations within s2m in SARS-CoV-2 emerging strains. For the Delta variant, we identified a G>U mutation at position 29,742 of the s2m (position 15 within the 41 nt of s2m, hence named here s2m G15U) that first appeared in January of 2020. The results of this analysis revealed a strong correlation between the s2m G15U and the Delta variant of SARS-CoV-2, highlighting a potential advantage conferred by the G15U mutation.

Recent evidence suggests that SARS-CoV-2 is evolving to evade the host immune response in vaccinated individuals, which makes studying the s2m and other structurally conserved elements crucial, in addition to the S protein (Andrews et al. 2022; Farinholt et al. 2021). Thus, in this work, we studied the dimerization properties of the s2m G15U and analyzed its interactions with the human miR-1307-3p, further expanding our breadth of knowledge surrounding s2m (Chan et al. 2020; Alam and Lipovich 2021). Better understanding of the role of conserved RNA structures and how interactions with host RNAs mediate their function will allow us to be better prepared for future variants, especially when considering the increased transmissibility and illness severity caused by the prevalent SARS-CoV-2 Delta variant (Luo et al. 2021; Liu and Rocklöv 2021).

## Results and Discussion

### The SARS-CoV-2 Delta variant s2m acquired the G>U mutation at position 15

We performed a bioinformatics analysis to monitor sequence mutants of the SARS-CoV-2 s2m in cases worldwide, specifically targeting India, the United Kingdom, and the United States, three countries which reported a significant number of SARS-CoV-2 sequences to the GISAID database (Elbe and Buckland-Merrett 2017). For this mutational analysis we used a custom R script that extracted and aligned the s2m sequences in all collected SARS-CoV-2 cases (Frye et al. 2023). Our bioinformatics analysis, which sampled SARS-CoV-2 sequences from April 2020 to October 2021, identified a G to U mutation at position 15 of the s2m, denoted as s2m G15U. The earliest observation of this s2m mutation was in April of 2020, within the United Kingdom and United States, at 0.23% prevalence. While the s2m G15U prevalence remained low for the majority of 2020, we observed an increase in s2m G15U prevalence in April 2021 and May 2021 in the United Kingdom and United States, respectively (Figure 3A, orange and blue, dashed lines). Interestingly, this increase was concurrent with the rise of the SARS-CoV-2 Delta variant (PANGO lineages B.1.617.2 + AY.*) (Figure 3A, orange and blue, solid lines). The presence of the s2m G15U rose from 0.23% in March 2021 to a 95.97% plateau in June 2021 in the United Kingdom, and from 0.82% in April of 2021 to a 92.40% plateau in July 2021 in the United States. This was simultaneous with the increase in the Delta variant, which rose from 2.03% in March 2021 to 95.68% in June 2021 in the United Kingdom, and from 0.62% in April of 2021 to 92.86% in July 2021 in the United States. We determined that the s2m G15U and Delta prevalence maintained an average Pearson’s correlation coefficient (R) of 0.9999 for both the United Kingdom and United States from April 2020 through October 2021, spanning the range of our entire bioinformatics analysis (Supplemental Figure 1A and 1B). Additionally, as a control, we studied the Alpha variant (PANGO lineages B.1.1.7 + Q.*) of SARS-CoV-2 and found that only 0.30% of Alpha sequences from November 2020 to August 2021, when this variant was dominant, contained the s2m G15U (Supplemental Figure 2). This shows that the s2m G15U mutant, while occurring in Alpha at low prevalence, did not rise in prevalence as it had in the Delta variant.

**Figure 3:**
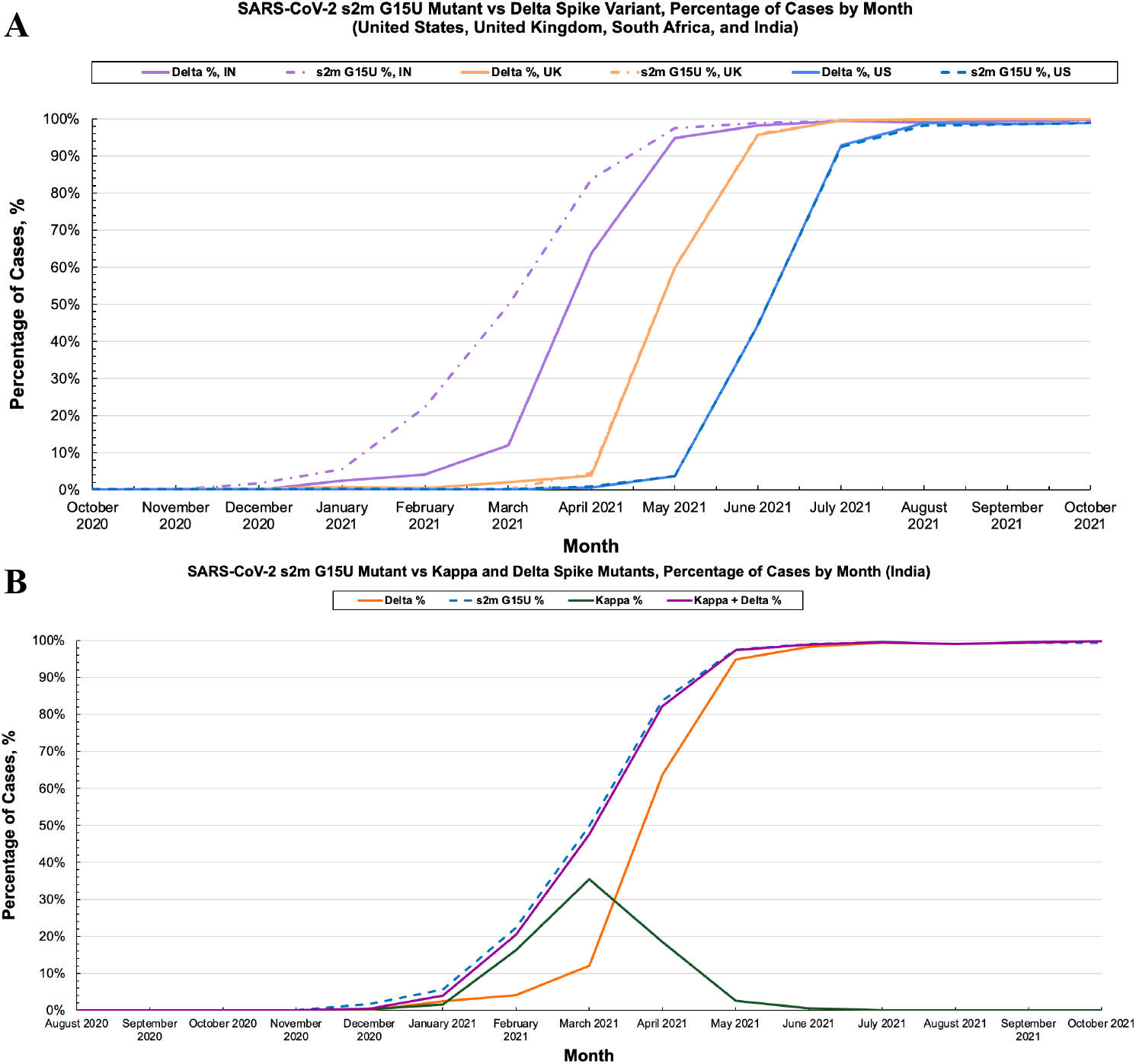
**A)** Prevalence of the s2m G15U mutation (dashed) and Delta variant (solid) within total reported SARS-CoV-2 cases in the United States (blue), United Kingdom (orange), and India (purple). **B)** The s2m G15U mutant (blue dashed line) rose in prevalence before the rise of the Delta variant (orange solid line). Kappa (green solid line) was also found to retain the s2m G15U mutation and comprised most cases within India that were not accounted for with Delta. Combination of the Kappa and Delta variants prevalence is tightly correlated with the s2m G15U mutant, mirroring trends later observed as Delta spread worldwide.

While the s2m G15U mutation and Delta variant show a strong positive correlation in prevalence throughout the spread of Delta worldwide, we noted a chronological discrepancy in the s2m G15U and Delta prevalence in India from November 2020 to March 2021, in which the s2m G15U had a noticeable difference in correlation from Delta (Figure 3A, purple dashed and solid lines). This difference is of particular interest given that the Delta variant originated in India. Here, the s2m G15U mutant increased significantly from 1.74% in December 2020 to 97.46% by May 2021, with the largest deviation between s2m G15U and Delta in March 2021, as the Delta variant comprised of only 12.06% of submitted cases compared to the 49.87% of s2m G15U. However, through May of 2021, both data sets converged as they reached 94.86% and 97.46% for Delta and s2m G15U, respectively. Calculation of this data set revealed R= 0.96619 (Supplemental Figure 1C, green), indicating a positive correlation with greater deviation compared to prior data sets. In efforts to determine the cause of this discrepancy between s2m G15U and Delta, we screened all Delta cases for the percentage of s2m G15U containing sequences and found that the Delta variant was comprised of 99.32% s2m G15U (Supplemental Figure 3, blue dashed line). We also analyzed India for other dominant VOCs which may contain the s2m G15U and noted that the Delta variant arose concurrently with a sister variant, Kappa (PANGO lineage B.1.617.1), and found that the s2m G15U is present in 99.05% of all submitted Kappa cases (Supplemental Figure 3). This finding indicated that Kappa had accounted for a majority of the s2m G15U cases that are not observed within Delta sequences. Combination of the Kappa and Delta prevalence, when plotted against s2m G15U, revealed R= 0.9997 (Supplemental Figure 1C, purple), indicating a strong positive correlation between the rise in s2m G15U and overall combined incidence of the Kappa and Delta variants. Thus, we attributed the rise of the s2m G15U to a concurrent increase in Kappa and Delta cases. It is to be noted, however, that the Kappa variant was subsequently outcompeted by Delta in March of 2021 (Figure 3B, green and orange), attributed to Delta’s increased ACE2 binding affinity and antibody escape (Liu et al. 2022; Kannan et al. 2021; Tian et al. 2021). This trend continues following the drop in Kappa prevalence, in which the combined Kappa and Delta variant prevalence maintains its high correlation with the s2m G15U mutant (Figure 3B, blue dashed line and purple solid line).

The strong positive correlation between the s2m G15U and the Delta variant through its rise in India, and eventual spread worldwide, indicates that the s2m G15U rose in tandem with Delta and was found to be in an average of 99.32% of all Delta sequences. Thus, given the presence of s2m G15U in such vast number of Delta SARS-CoV-2 cases and the high conservancy and predicted roles of the s2m in the viral life cycle, this warranted its further characterization.

### S2m G15U secondary structure characterization

To evaluate the effect of the G15U mutation on the secondary structure of the s2m, we utilized ^1^H NMR spectroscopy, comparing the imino proton resonance region for the reference s2m and s2m G15U at 19°C. The reference SARS-CoV-2 s2m imino proton resonances were previously assigned and thus served as a baseline for our work (Imperatore et al. 2022; Wacker et al. 2020). The previously assigned imino protons corresponding to base pairs in the s2m lower stem (13.19 ppm - G37, 13.42 ppm - U38 and 11.62 ppm - G39) are identical between the two spectra (Figure 1B). This indicates that the s2m G15U lower stem is not perturbed, consistent with the predicted secondary structures of reference s2m and s2m G15U (Figure 1A). Similarly, the resonance previously assigned to the U31 imino proton has the same chemical shift (13.93 ppm) in the spectrum of reference s2m and s2m G15U. We used ^1^H-^1^H NOESY NMR spectroscopy on s2m G15U at 19°C to confirm these assignments and assign the remaining imino proton resonances which differ in chemical shift between the reference s2m and s2m G15U. As expected, the resonances assigned to U31 and U38 imino protons (13.93 ppm and 13.42 ppm) have a strong NOE with the H2 of the corresponding adenines they are base paired with (A13 H2 at 7.23 ppm for U31 NH and A4 H2 at 7.18 ppm for U38 NH) (Figure 1C). In addition, the resonance at 13.66 ppm gives a single strong NOE at 7.39 ppm, a signature of an A-U base pair, allowing us to assign it to the U15 imino proton (Figure 1C). This result confirms that the G15U mutation closes the base pair U15-A29, as predicted for the s2m G15U secondary structure (Figure 1A, top). Based on its neighboring base pairs, the U31 imino should give rise to NOEs with the G30 and G32 imino protons, and indeed, two cross peaks are observable, at 13.5 ppm and at 11.76 ppm, for the U31 imino proton resonance at 13.93 ppm. Based on its neighboring base pairs, the U15 imino proton should give rise to two NOEs: with the imino protons of G30 and G28. We only observe one clear NOE at 13.5 ppm from the U15 imino proton at 13.66 ppm. Since the imino at 13.5 ppm gives NOEs to both the U31 imino and the U15 imino, we unambiguously assign it to the G30 imino, as this is the only imino proton in the proximity of both U31 and U15 imino protons. Then, by exclusion, the second NOE observed for the U31 imino (13.93 ppm; 11.76 ppm) leads to the assignment of the resonance at 11.76 ppm to the G32 imino proton. This assignment is also supported by the observed NOE between the G32 imino at 11.76 ppm and the A13 H2 at 7.23 ppm. We also observed NOEs between the imino protons of U38 and G37 (13.42 ppm; 13.19 ppm), as well as between the imino of G37 and the A4 H2 at 7.18 ppm, confirming their previous assignment (Figure 1D) (Imperatore et al. 2022; Wacker et al. 2020). A weak NOE is also observed between the imino of G39 at 11.62 ppm and the A4 H2 at 7.18 ppm, confirming the G39 imino assignment (Figure 1C). Finally, there are two remaining imino proton resonances at 12.03 ppm and 12.7 ppm, which do not give rise to NOEs that would allow us to unambiguously assign them to G28 and G17 iminos. The imino proton resonances of U1 and U2 are not visible in either spectrum due to the dynamic nature of these end base pairs (Imperatore et al. 2022). Based on these assignments we conclude that aside from the obvious mutation which closes the U15-A29 base pair, the s2m G15U secondary structure does not deviate far from that of the reference s2m.

To further characterize the effect this newly formed base pair has on the stability of the s2m, we performed UV thermal denaturation experiments on s2m G15U and reference s2m, monitoring the absorbance changes at 260 nm (Figure 4). The melting temperature, T_m_, of each structure was determined from the first derivative plots (Supplemental Figure 4) to be 44°C and 47°C for the reference s2m and s2m G15U, respectively. This indicated that, as expected, the closing of the U15-A29 base pair in the s2m G15U upper stem increases the overall stability of the s2m G15U (Figures 2 and 4), which may have implications for the virus’s ability to dimerize and form extended duplex structures. In our previous work, we established that the s2m elements of SARS-CoV and SARS-CoV-2 have different dimerization abilities and determined that this is due to the presence of a single nucleotide mutation in s2m G31U located in the s2m upper stem (Imperatore et al. 2022). Such dimerization events in viruses have been shown as critical for genome packaging, inducing conformational changes for translational regulation, reverse transcription processes, as well as being important for genetic recombination (Tengs et al. 2013; Terada et al. 2014; Berkhout and van Wamel 1996; Aagaard et al. 2004; Mikkelsen et al. 2000; Moore et al. 2009). Therefore, next we evaluated the effect of the single nucleotide mutation, the G15U, on the Delta SARS-CoV-2 s2m dimerization.

**Figure 4.**
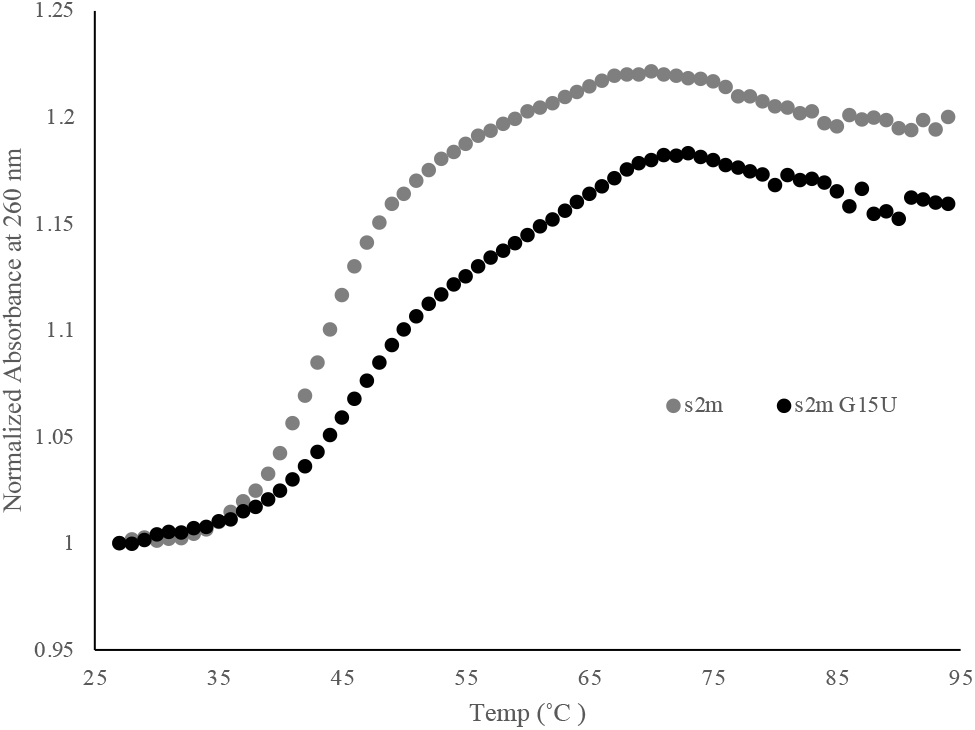
UV thermal denaturation of the reference s2m and s2m G15U at 1 μM in 10 mM cacodylic acid pH 6.5 and in the absence of MgCl_2_. The melting temperatures, T_m_, were 44°C and 47°C for the reference s2m and s2m G15U, respectively. The data points were normalized using the first temperature as reference.

### Characterization of the SARS-CoV-2 s2m G15U dimerization

Our previous work established the presence of a 4 nt palindromic “GUAC” sequence within the terminal loop of the s2m element that participates in loop-loop kissing dimerization (Imperatore et al. 2022). These kissing dimer structures are formed through base pairing between two RNA hairpin loops that contain complementary unpaired nucleotides. Kissing dimers forming with palindromic sequences have been previously reported in other viral systems to be essential for template switching and recombination, a process important for repairing genetic mistakes and increasing diversity within the virus-which is critical for the evolution of new viral strains (Andersen et al. 2003; Mikkelsen et al. 2000). These kissing dimers can then be converted to a thermodynamically stable extended duplex structure through the chaperone activity of the capsid protein when in the context of the entire viral genome (Figure 2). However, *in vitro*, short isolated kissing dimers have been reported to form extended duplex conformations spontaneously, such as in hepatitis C virus (HCV) and human immunodeficiency virus (HIV-1) (Muriaux et al. 1996; Shetty et al. 2010; Mihailescu and Marino 2004). In the case of HIV-1, recombination events have led to high rates of immune escape and drug resistance. Thus, given the potential of s2m to be involved in recombination events it is important to determine if the G15U mutation affects its dimerization, as SARS-CoV-2 is still developing new variants (Morris et al. 1999; Rambaut et al. 2004; Dubois et al. 2018).

Therefore, to analyze the G15U mutation’s effect upon dimerization of the SARS-CoV-2 s2m, we used TBE/TBM nondenaturing gel electrophoresis, a method used previously in various viral systems to distinguish between kissing dimer versus duplex conformations of dimer initiation sites (Sun et al. 2007; Berkhout et al. 2002; Li et al. 2006; Kenyon et al. 2013). Conditions which maintain MgCl_2_ concentrations (TBM) stabilize the formation of the kissing dimer; however, upon chelation of the Mg^2+^ ions by EDTA in TBE, the kissing dimer dissociates, whereas the duplex conformation, which do not require Mg^2+^ ions for stability, is retained. When MgCl_2_ is retained in the TBM gel, a prominent monomer band is present for both the reference s2m and s2m G15U (Figure 5, right, arrows 1 and 1’). There are two dimer bands present for the reference s2m (Figure 5, right, arrows 2 and 3), and a single faint dimer band is present for s2m G15U (Figure 5, right, arrow 3). Our previous report has established that these two upper bands in the reference s2m (Figure 5, right, arrows 2 and 3) correspond to a kissing dimer and extended duplex, respectively (Imperatore et al. 2022). Although both the kissing dimer and extended duplex conformations contain the same number of nucleotides, their migration patterns can differ in native gels as these separate RNAs based on both size and shape. Duplex structures are linear, whereas kissing dimers can be present in either a bent or linear state. This would give rise to a different shape that may affect the migration pattern, explaining the difference in the two bands (Rist and Marino 2002; Ennifar et al. 1999). Although unlikely in the context of the entire genome, when the reference s2m is isolated in its 41 nt state, spontaneous conversion to an extended duplex structure is possible and has been reported in studies involving isolated dimerization initiation sites in other viruses (Muriaux et al. 1996; Shetty et al. 2010; Mihailescu and Marino 2004). Similarly, it is interesting to note that, despite having the same size, the monomer bands for the reference s2m and s2m G15U migrate differently in the TBM gel (Figure 5, right, arrow 1 and 1’), suggesting that they have different monomer conformations in the presence of MgCl_2_. Our MD simulations for the reference s2m revealed that it has a kinked 3D structure; however, it is possible that the additional U15-A29 base pair in the upper stem of s2m G15U results in a more linear structure that will migrate further through the TBM gel and MD simulations are in progress in our laboratories to test this hypothesis (Kensinger et al. 2022). Thus, we cannot determine if the single dimer band observed for the s2m G15U in the TBM gel (Figure 5, right, arrow 3) is originating from a mixture of linear kissing dimer and duplex conformations which will migrate at the same position, or from the duplex conformation formed spontaneously from an unstable kissing dimer during incubation and migration through the gel (Shetty et al. 2010).

**Figure 5.**
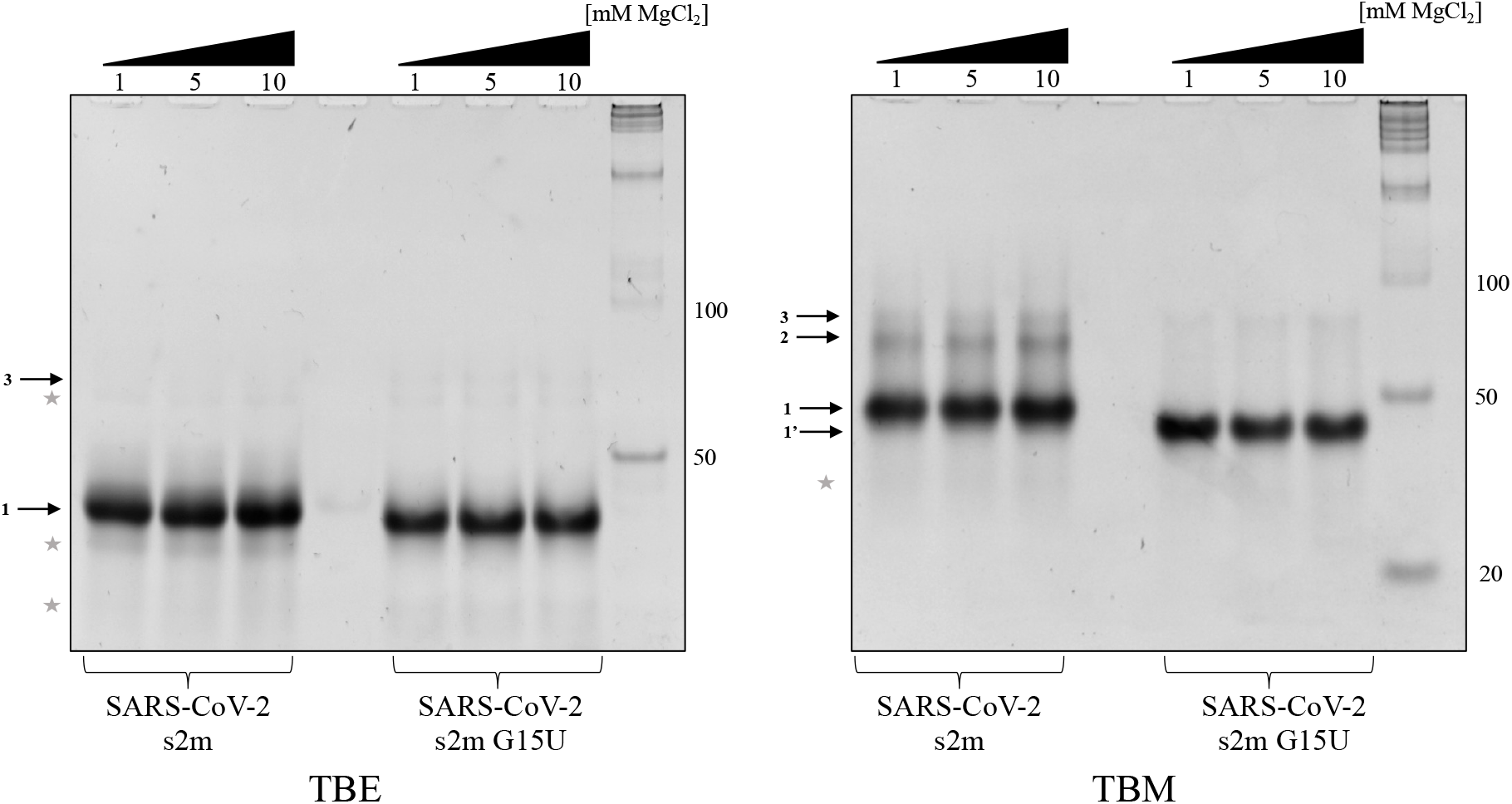
TBE (left) and TBM (right) gel non-denaturing electrophoresis analysis of the reference SARS-CoV-2 s2m and the s2m G15U. The two s2m elements were compared in the presence of Mg^2+^ where 1 μM RNA was incubated in the presence of 1, 5, or 10 mM MgCl_2_ for 1 h then split between two gels-TBE and TBM. In the TBE both s2m elements prefer their monomeric states (left, arrow 1). In TBM, the monomeric states are still preferred (right, arrows 1 and 1’). Different migratory patterns may be observed due to different shapes of the s2ms, as described in the text. Arrows 2 and 3 indicate the kissing dimer and extended duplex, respectively. Degradation products are noted with *.

In the TBE gel, after an hour incubation with MgCl_2_, a prominent monomer s2m band exists for both the reference s2m and the s2m G15U mutant (Figure 5, left, arrow 1). We also observed faint monomer bands marked by * originating from degradation products. There are also faint dimer bands present in the TBE gel, which likely originate from the duplex conformations of the intact s2m elements and of its minor degradation products (see below). It is important to note that the TBM gel was run for 4 hours in the presence of MgCl_2_, as opposed to the TBE gel that was run for 2 h, allowing more time for spontaneous conversion of the s2m kissing dimer to the duplex conformation, and hence, there is a more prominent duplex band in the TBM gel (Figure 5, right, arrow 3), as compared to the TBE gel.

However, in the context of the full virus, this conversion from the kissing dimer to the extended duplex conformation would not occur spontaneously and would instead require the assistance of the N protein’s chaperone activity to facilitate the transition. This process is dependent on the overall stability of the s2m element as base pairs in each monomer will have to be broken, disrupting the monomeric hairpin structure. In coronaviruses, the N protein is responsible for packaging the RNA into ribonucleoprotein complexes to form the mature virion prior to being exported from the host cell (Aduri et al. 2013; Zúñiga et al. 2007). Our previous work has shown that the SARS-CoV-2 N protein selectively acts as a molecular chaperone to convert the reference s2m kissing dimer structure to the stable duplex by a similar mechanism observed for the dimer initiation sites of HIV-1 and HCV (Shetty et al. 2010; Imperatore et al. 2022; Rist and Marino 2002; Shimakami et al. 2012). In HIV-1, a retrovirus, dimerization of the RNA is important for the packaging of two copies of the genome in the mature virion (Dubois et al. 2018). In HCV, a single-stranded RNA virus, this dimerization is suggested to be a regulator of RNA replication and viral genome packaging (Shetty et al. 2010). Additionally, the dimer initiation sites in HIV-1, HCV, and in the murine leukemia virus have been proposed to be implicated in recombination events (Masante et al. 2015; Greatorex 2004; Torrent et al. 1994). We proposed previously that the s2m dimerization mediated by the viral N protein may facilitate SARS-CoV-2 recombination; thus, here we investigated if the G15U mutation affects the s2m dimerization properties (Imperatore et al. 2022).

To test the chaperone ability of the N protein to convert the s2m G15U to the duplex structure, as compared to the reference s2m, a 1 μM s2m RNA sample was incubated with 1 mM MgCl_2_ for 30 minutes, followed by the addition of 2 μM N protein for an additional 30 minutes. The SARS-CoV-2 main protease (P) protein and bovine serum albumin (BSA) were used as control proteins. After the incubation with the N or control proteins, proteinase K was added to degrade the proteins prior to loading the samples on the TBE and TBM gels. As demonstrated previously, in TBM both the reference s2m and s2m G15U preferred their monomeric states (Figure 6, right, arrows 1 and 1’) in the absence of the N protein; however, in the presence of the N protein the two upper bands in the reference s2m became more prominent (Figure 6, right, arrows 2 and 3) (Imperatore et al. 2022). Similarly, the dimer band of s2m G15U, which was barely visible on its own, becomes clear in the presence of the N protein (Figure 6, right, arrow 3). As discussed above, we cannot distinguish if the s2m G15U kissing dimer is unstable (hence the dimer band indicated by arrow 2 is absent in s2m G15U) or if it is a linear structure that migrates at the same position with the duplex conformation. When the Mg^2+^ ions are chelated in the TBE gel, a prominent dimer band remains for both reference s2m and s2m G15U that were incubated with the N protein (Figure 6, left, arrow 3), indicating that the N protein facilitates the conversion to the duplex conformation in both samples. Note that the faint dimer bands marked by * do not line up with the main duplex bands (Figure 6, left, arrow 3), indicating that they originate from dimerization of the degradation products. The addition of the P protein or BSA to the sample had no effect, validating the specificity of the N protein as a molecular chaperone.

**Figure 6.**
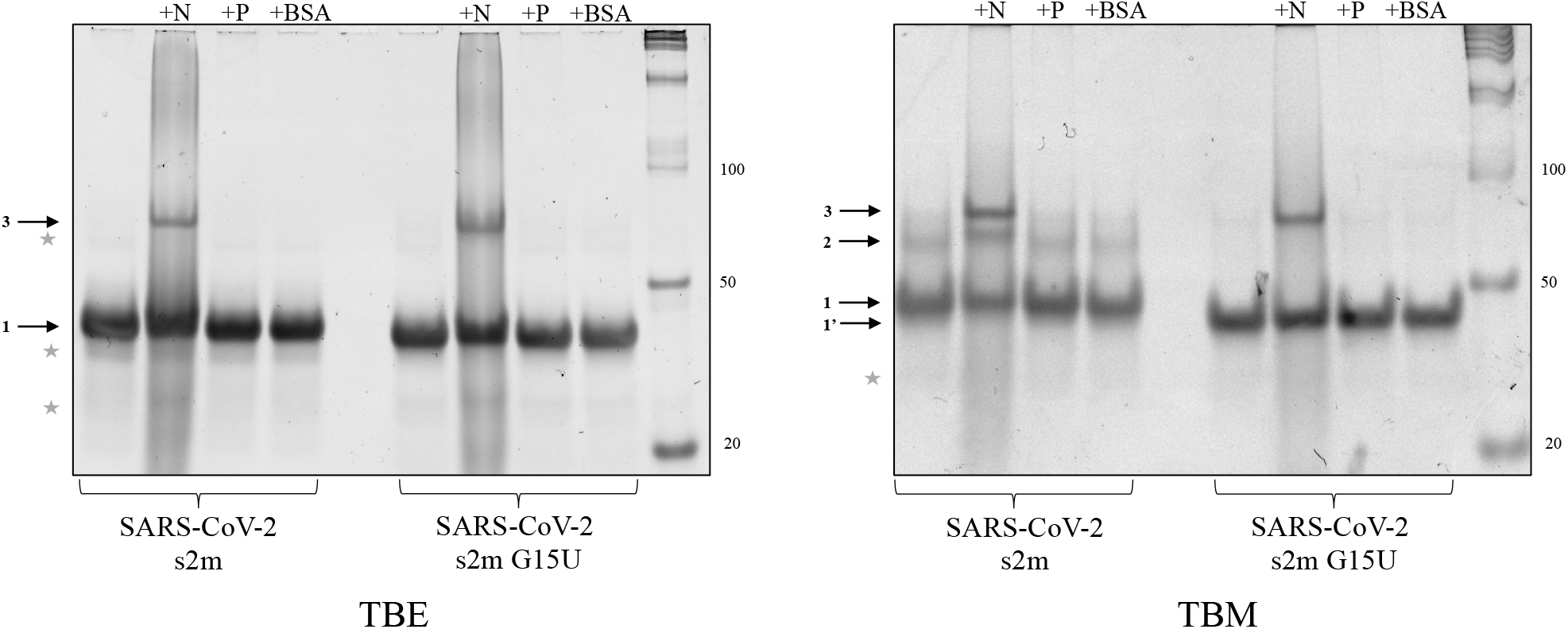
TBE and TBM non-denaturing gel electrophoresis of the reference s2m and s2m G15U converted to an extended duplex structure by the viral N protein. Both s2m elements were incubated with 1 μM MgCl_2_ for 30 min, an additional 30 min with the N protein, and 5 min with proteinase K to digest the proteins prior to the samples being split in TBE (left) and TBM (right) conditions for electrophoresis. In both cases, the s2m remains preferentially monomeric (arrow 1 and 1’) in the absence of the N protein. The N protein converted both elements to an extended duplex structure (arrow 3), and in the TBM gel, the N protein also stabilized the kissing dimer of the reference s2m (right, arrow 2). However, the band corresponding to the kissing dimer of the s2m G15U sample is still not observed. The P protein and BSA were used as controls and had no effect on the conversion of either s2m kissing dimer to the duplex conformation.

Based on these results, we conclude that while the reference s2m and s2m G15U are converted by the N protein to the duplex conformation, the s2m G15U kissing dimer conformation is either not as stable or forms a different conformation than the reference s2m kissing dimer. Specifically, the absence of an s2m G15U kissing dimer band at the same position observed for the reference s2m kissing dimer (Figure 5, right arrow 2) indicates that the s2m G15U kissing dimer could be linear, migrating at the same position with the linear duplex conformation, different than the kissing dimer structure of the reference s2m. Alternately, this could indicate that the s2m G15U kissing dimer is able to convert spontaneously to the duplex conformation in the absence of the N protein more efficiently than the reference s2m; however, considering that the s2m G15U upper stem is more stable and must be opened during this conversion, this scenario is less likely. A difference in kissing dimer conformation is more probable as the monomers of these two s2m elements have a different shape based on their different migration in the TBM gel (Figure 5 right, arrows 1 and 1’). Despite these observed differences in the s2m G15U kissing dimer versus the reference s2m kissing dimer, we further demonstrate that this mutation does not affect the ability of the viral N protein to convert the s2m G15U kissing dimer to a stable duplex structure, which is the *in vivo* process of duplex formation. It has been suggested that the N protein preferentially binds to double stranded RNA segments in the central region of the s2m, specifically nucleotides G28, G30, and U31, making a stable protein-RNA complex for virion packaging (Padroni et al. 2022; Morse et al. 2023). Therefore, the closing of the U15-A29 base pair, despite making the s2m G15U upper stem more stable, which would be unfavorable for the conversion of the s2m G15U kissing dimer to the duplex conformation, may compensate for this unfavorable effect by further assisting in N protein binding.

The findings that the G15U mutation does not affect the ability of the N protein to convert the s2m G15U kissing dimer to the duplex conformation are contrasting the effect another single nucleotide mutation in the s2m upper stem (G31U) has on this conversion. In our previous work, we showed that the G31U mutation that exists between the SARS-CoV and SARS-CoV-2 s2m was responsible for drastic differences in the s2m dimerization, even in the presence of the N protein (Imperatore et al. 2022). The SARS-CoV s2m readily forms kissing dimers that could not be converted to the extended duplex by the N protein. Conversely, the SARS-CoV-2 reference s2m which differs by only two nucleotides (only one of which, G31U, we showed is responsible for the dimerization differences) from SARS-CoV s2m, exists primarily as a monomer that forms kissing dimers which are subsequently converted to the duplex conformation by the N protein (Imperatore et al. 2022). While the single nucleotide G31U mutation was shown to be responsible for this difference in s2m dimerization properties between SARS-CoV s2m and the reference SARS-CoV-2 s2m, we show here that the s2m G15U single nucleotide mutation present in the Delta variant has limited effects on the element’s dimerization (Imperatore et al. 2022).

### SARS-CoV-2 s2m G15U interactions with miR-1307-3p

We and others previously identified two miR-1307-3p binding sites on the s2m element in the reference SARS-CoV-2 3’-UTR, and we revealed its specificity for this miR (Imperatore et al. 2022; Arisan et al. 2020; Balmeh et al. 2020; Chan et al. 2020; Alam and Lipovich 2021). The exact role of miR-1307-3p in the SARS-CoV-2 infection is not known; however, it has been shown that miR-1307-3p is the highest expressed microRNA in Vero cells infected by SARS-CoV-2 and the levels of miR-1307-3p are elevated in severe cases of COVID-19 as compared with mild cases (Zarei Ghobadi et al. 2022; Arisan et al. 2022). Thus, we investigated if the G15U mutation within the s2m has any effect upon its interactions with miR-1307-3p. The binding interactions between miR-1307-3p and s2m G15U as compared to the reference s2m were assessed by TBE/TBM native PAGE. The reference s2m and s2m G15U were boiled and snap cooled, then incubated with 1 mM MgCl_2_ for 30 mins, after which miR-1307-3p was added in 1:1 and 1:2 s2m:miR ratios and incubated for an additional 30 mins at room temperature. The samples were then split and electrophoresed in TBE and TBM conditions, respectively. In the TBE gel, two bands are present for miR-1307-3p, a monomer and dimer (Figure 7, left, arrows 1 and 3), while both the reference s2m and s2m G15U are present as monomers (Figure 7, left, arrow 2). As miR-1307-3p was added to the s2m samples, two upper bands are present for the reference s2m, which were assigned to the binding of one miR-1307-3p molecule (Figure 7, left, arrow 4’) and of two miR-1307-3p molecules, respectively (Figure 7, left, arrow 5) (Imperatore et al. 2022). However, these upper bands are barely visible for s2m G15U, indicating that miR-1307-3p has reduced affinity for this mutant. Note that there is a faint band at almost the same position indicated by arrow 5 in the free reference s2m and s2m G15U which corresponds to the duplex conformation (82 nt), which is very close in size to the complex formed by s2m with two miR-1307-3p molecules (85 nt). When the Mg^2+^ ions are retained in the TBM gel, the complex of the reference s2m with two miR-1307-3p molecules is stabilized (Figure 7, right, arrow 5) and a new higher order complex appears which we assigned to an s2m dimer interacting with two miR-1307-3p molecules, as described in our previous work (Figure 7, right, arrow 6) (Imperatore et al. 2022). In contrast, while Mg^2+^ ions stabilize the complex formed by s2m G15U with two miR-1307-3p molecules (Figure 7, right, arrow 5), the band intensity corresponding to these complexes is very low. This result suggests that the closing of the U15-A29 base pair creates a more stable s2m upper stem, making it harder for the miR-1307-3p to invade it, and ultimately lowering its binding affinity.

**Figure 7.**
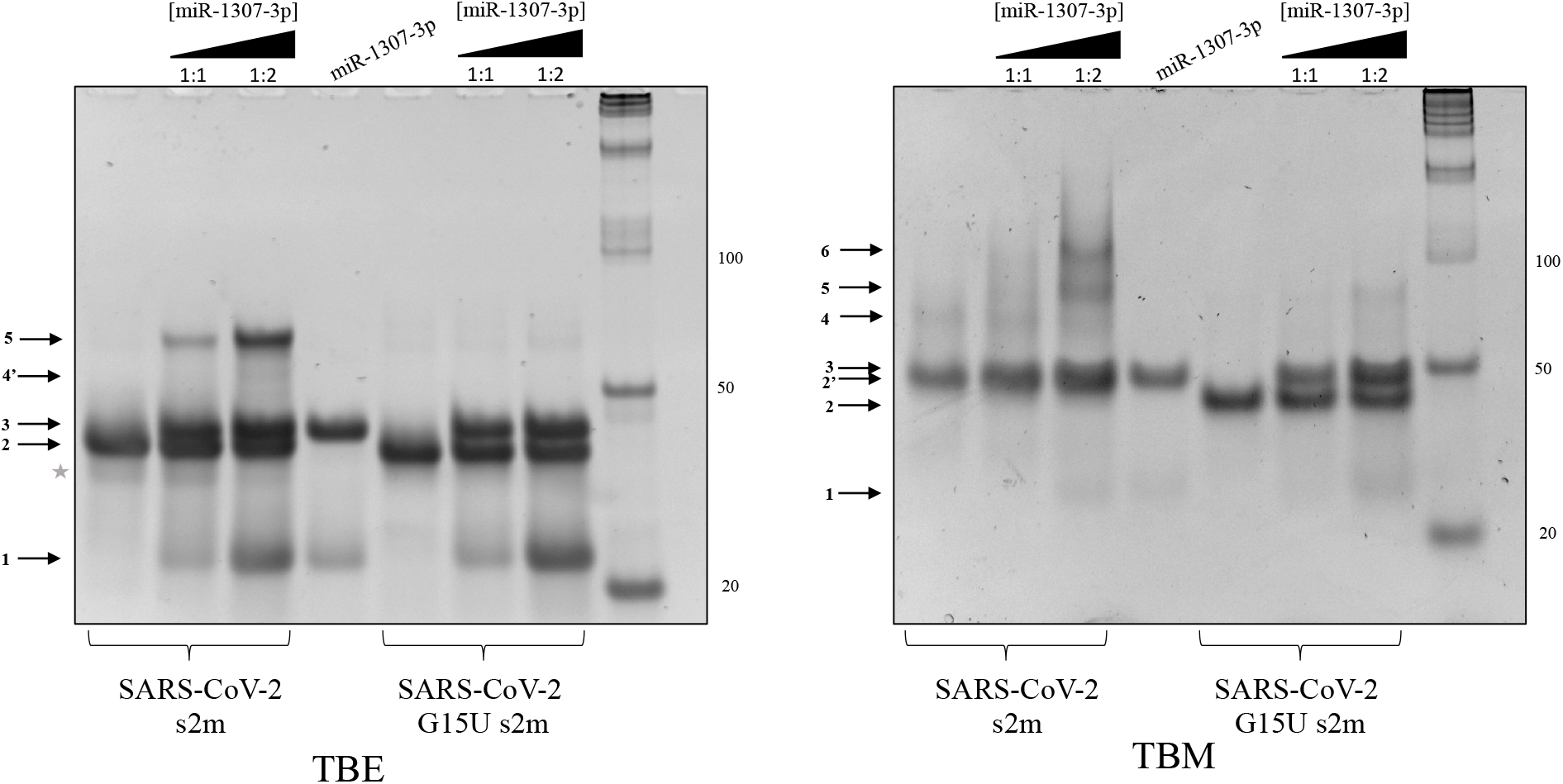
Non-denaturing TBE (left) and TBM (right) gels of s2m and s2m G15U binding to miR-1307-3p in 1:1 and 1:2 s2m:miR ratios in the presence of 1 mM MgCl_2_. In TBE, both s2m elements prefer their monomeric states (arrow 2) whereas the miR-1307-3p exists as a monomer and dimer (arrows 1 and 3). As miR is added, complexes are formed between the two (arrows 4’ and 5). The band intensity of the complexes formed by s2m G15U with miR-1307-3p is drastically reduced. In TBM, the reference s2m forms a stable complex with two miR-1307-3p (arrow 5) and a complex formed by two s2m each bound to one miR-1307-3p is also apparent (arrow 6) (Imperatore et al. 2022). Arrow 4 shows the kissing dimer formed by the reference s2m. s2m G15U complex with miR-1307-3p is stabilized in the presence of Mg^2+^(arrow 5). The monomers of s2m and s2m G15U migrate differently (arrows 2 and 2’), as described in the text.

Disease severity in patients with SARS-CoV-2 has fluctuated throughout the pandemic, ranging from asymptomatic to mild to severe. In severe cases of COVID-19, hyperactive immune responses lead to the overproduction and release of cytokines, termed the “cytokine storm.” These cytokines are proteins critical for the regulation of the host’s defense mechanisms (Vial and Descotes 1995). An example of cytokines includes interleukins (ILs), such as IL18, and their receptors (e.g., IL6R, IL10RA, IL10RB, IL12RB2, IL17RA), and it has been shown that their overexpression is linked to the onset of cytokine storms in COVID-19 patients (Agarwal et al. 2015). A suggested regulator of these ILs and their receptors is host miR-1307-3p (Imperatore et al. 2022; Mehta et al. 2020; Yang et al. 2020). MiR-1307-3p has also been proposed to reduce glucose regulating protein 78 (GRP78), a protein thought to aid in binding of the SARS-CoV-2 S protein to the host cell’s receptors (Ibrahim et al. 2020; Elfiky 2020; Carlos et al. 2021). By allowing the virus to hijack this miR, GRP78 abundance in the cells would be increased, aiding viral entry into the cell. Viruses have been shown to hijack host miRNAs for their own benefit (Trobaugh and Klimstra 2017; Guo and Steitz 2014). One example of this is HCV, which uses miR-122 to enhance its replication (Shimakami et al. 2012; Jopling et al. 2005). Previously, upon finding that the reference s2m binds two copies of miR-1307-3p we proposed that this miR is hijacked by SARS-CoV-2 for the benefit of the virus (Imperatore et al. 2022). However, considering these new results, our previous hypothesis has to be amended as it seems that the virus has evolved to reduce its interactions with this miR in the Delta variant. We show that the s2m G15U has lower affinity for miR-1307-3p as compared to the reference s2m and suggest the newly formed U15-A29 base pair formation as causative for these diminished interactions, as the s2m upper stem must be opened for new base pairs to be formed with miR-1307-3p. This implies that miR-1307-3p may instead have a detrimental effect on the virus, as our data supports that the virus preferentially evolved to reduce this interaction. Recent work has identified that host miRs may target the viral genome as part of the immune response, in which case, the virus would preferentially evolve to evade this attack (Skalsky and Cullen 2010; Huang et al. 2007; Lecellier et al. 2005; Otsuka et al. 2007). MiRs normally bind to the target mRNA 3’-UTR to signal its degradation and/or to regulate its translation (O’Brien et al. 2018). Often, the miR binding sites on mammalian mRNAs are highly conserved, preserving the regulatory role of these miRs (Friedman et al. 2009). Given the high conservation of the s2m in the 3’-UTR of several viral families, it is possible that the host immune response could employ miRs to target this element, and several miRs are dysregulated upon SARS-CoV-2 infection (Farr et al. 2021). Thus, by targeting the s2m of SARS-CoV-2, the host may be utilizing miR-1307-3p to alter the translation of viral proteins (Panda et al. 2022). It is interesting to note that since the SARS-CoV-2 subgenomic RNAs share the same 3’-UTR with the genomic RNA, miR-1307-3p could target the translation of both the viral structural and nonstructural proteins. The host miR-1307-3p has been shown to be upregulated in SARS-CoV-2 infected cells, up to a 38-fold increase, and is implicated in the silencing of SARS-CoV-2 post transcriptionally (Panda et al. 2022; Zarei Ghobadi et al. 2022; Arisan et al. 2022). Additionally, increased expression of miR-1307-3p has been shown to result in decreased replication of the SARS-CoV-2 genome (Zarei Ghobadi et al. 2022; Soremekun et al. 2020). Therefore, mutations which decrease the binding interactions of miR-1307-3p with the s2m, such as those we observe in the s2m G15U mutant, would be optimal for overall viral fitness of Delta compared to the previous variants that contain the reference s2m.

Interestingly, our recent work has shown that the Omicron variant, the most prominent viral strain to emerge after Delta, contains a 26 nt deletion within the s2m element that eliminates the miR-1307-3p binding sites (Frye et al. 2023; Colson et al. 2022). Omicron has also been shown to have further reduced disease severity and increased infectivity when compared to Delta and other prior variants (Chaguza et al. 2022). We emphasize that our results could be evident of a stepwise reduction of the miR-1307-3p binding sites as the s2m has evolved to reduce miR-1307-3p binding in Delta and subsequently removed the binding sites with the 26 nt deletion in Omicron (Frye et al. 2023). This trend supports the notion that SARS-CoV-2 is evolving towards reduced miR-1307-3p binding. Despite the lack of knowledge regarding this interaction, we suspect that if miR-1307-3p is in fact a crucial player in the antiviral immune response, the Omicron variant of SARS-CoV-2 greatly benefits through an overall reduced miR-1307-3p binding. Further, if miR-1307-3p is involved in the “cytokine storm,” this variant also benefits through lower disease severity, which is seen in patients diagnosed with the Omicron variant. Sequence alignments of the SARS-CoV-2 variants to other coronaviruses, specifically human “cold-like” viruses, revealed that the Omicron deletion within the s2m is similar to those of the common cold (Arisan et al. 2022). This further supports that removal of this region, and the decreased (or lack of) ability to bind miR-1307-3p may be a similar evolutionary step taken by SARS-CoV-2 to prevent severe, and potentially deadly, symptoms in the host by improving evasion of the host immune response.

However, it is to be acknowledged that this data alone cannot prove directly if this s2m G15U mutant does indeed confer a selective advantage to SARS-CoV-2, or is a result of a hitchhiking effect, where the mutations within the S protein which conferred an advantage to the Delta variant happened on a genome that already had the s2m G15U mutation. Nonetheless, the findings that the Omicron variant, which is not a direct descendant of the Delta variant, acquired the 26 nt deletion that removed the miR-1307-3p binding sites suggests that the abolishment of the miR-1307-3p interactions with the genomic and subgenomic viral RNA is advantageous to the virus. Further research is needed to elucidate the details of the role of miR-1307-3p in the host immune response to SARS-CoV-2 infections and viral genome dimerization.

In summary, we showed here that the s2m G15U mutation is present in 99.32% of the SARS-CoV-2 Delta variant, and that while this mutation does not affect the s2m dimerization properties in the presence of the N protein, it does reduce its affinity for the host miR-1307-3p, potentially having implications on the host response to SARS-CoV-2 infection and viral evolution to evade this response.

## Materials and Methods

### Bioinformatics Analysis

SARS-CoV-2 genomes were retrieved from the GISAID EpiCoV platform for the period of April 2020 to October 2021, and to facilitate alignment, only complete and high coverage genomes were selected. The hCoV-19 Wuhan SARS-CoV-2 virus (GISAID: EPI_ISL_402123) was used as a reference genome (Wu et al. 2020). Analysis of the s2m was performed using a custom R script dependent on the packages *BiocManager*, *Biostrings*, *DECIPHER*, *hiReadsProcessor*, *adegenet*, *stringr*, and *ape*, which was recently used to monitor s2m sequence and identify mutations (Frye et al. 2023; Huber et al. 2015). To manipulate and align sequences, we used the *Biostrings* and *DECIPHER* packages, managed by *BiocManager*. We trimmed aligned sequences to the s2m with *hiReadsProcessor* and employed *adegenet*, *stringr*, and *ape* for further processing and identification of mutations within s2m. The SARS-CoV-2 reference sequence was converted to a FASTA file and added to each batch of sequences, which contained less than 10,000 sequences each. The first 29,000 nucleotides of each sequence were removed to reduce computational time, and the sequences were realigned at the 3’ end. After performing the alignment, the script identified sequences with single nucleotide insertions that disrupt mutation identification by introducing gaps in all other sequences in the alignment. These sequences were removed, and the remaining sequences were re-aligned. From a total of 2,034,376 sequences analyzed, 1,980 were found to have single nucleotide insertions and were removed from this analysis. The script then extracted the s2m motif from all sequences in the final alignment and identified mutations relative to the reference sequence. Mutations and corresponding information, including accession ID, geographic location, and collection date, were output in a CSV file. The sequence alignments were completed for each of the SARS-CoV-2 variants and each geographical location of interest. This data was then processed and used for identification of s2m mutation trends.

### RNA Samples

The s2m and miR RNA sequences used in this study (Table 1) were purchased from Dharmacon Inc. and resuspended in 10 mM cacodylic acid, pH 6.5.

**Table 1.**
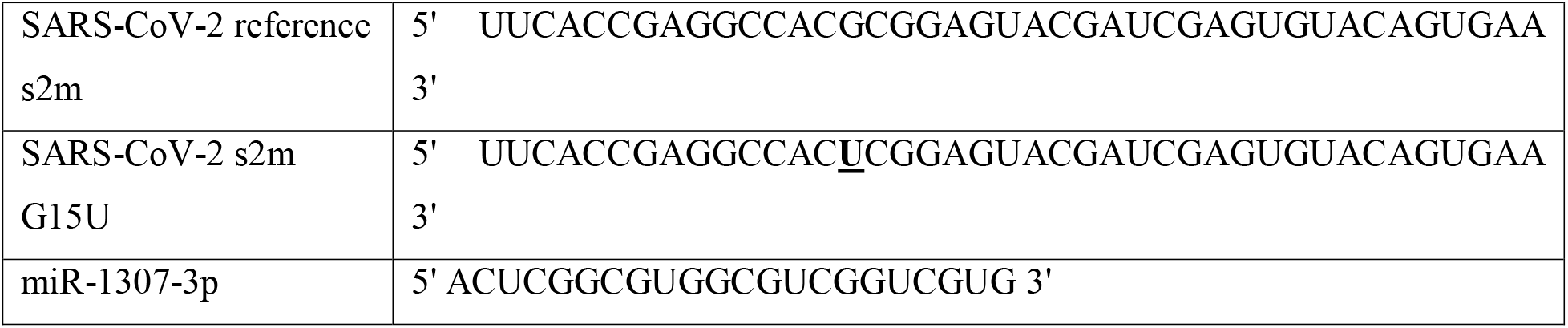
Oligonucleotide sequences used in this study.

### Nondenaturing Polyacrylamide Gel Electrophoresis

The RNA samples were diluted in ½x Tris-Boric Acid (TB) buffer to 1 μM, boiled for 5 min, then snap-cooled in dry ice and ethanol, after which, the samples were incubated at room temperature for 1 hour in the presence of varying concentrations between 1 mM and 10 mM of MgCl_2_. To evaluate the conversion of the kissing dimer to a thermodynamically stable extended duplex, 1 μM reference s2m and s2m G15U RNAs were incubated with 1 mM MgCl_2_ for 30 min followed by an additional 30-min incubation with 2 μM N protein (RayBiotech, Inc.). Prior to electrophoresing, proteinase K (New England Biolabs) was added for 10 min to digest the N protein. To analyze the interactions between the reference s2m and s2m G15U and miR-1307-3p, 1 μM s2m RNA was incubated with 1 mM MgCl_2_ for 30 min followed by and additional incubation with increasing concentration ratios of miR-1307-3p for 30 min. For all experiments, samples were evenly split after incubation and run in two separate gel conditions: Tris-Boric Acid EDTA (TBE) or Tris-Boric Acid with 5 mM MgCl_2_ (TBM). The samples were electrophoresed for 2 hours at 75 V in TBE and 4 hours in TBM using ½x TBE and ½x TBM running buffer, respectively. Visualization of the gels was done by staining in SYBR^®^ Gold cyanide dye and UV transillumination at 537 nm on an AlphaImager. All experiments were completed in triplicate.

### ^1^*H NMR Spectroscopy*

One-dimensional ^1^H NMR spectroscopy of the reference s2m and s2m G15U was performed at 19°C on a 500 MHz Bruker AVANCE NMR spectrometer equipped with TopSpin3.2 acquisition software. RNA samples at a concentration of 220 μM were prepared in 10 mM cacodylic acid, pH 6.5, in a 90:10 H_2_O:D_2_O ratio. Samples were boiled for 5 min and snap-cooled using dry ice and ethanol prior to data acquisition. Water suppression was carried out using the Watergate pulse sequence (Piotto et al. 1992).

A ^1^H-^1^H NOESY experiment with a 150 ms mixing time was conducted for s2m G15U at 19°C on a 500□MHz Bruker AVANCE spectrometer. The sample was prepared at 250 μM RNA in 10 mM cacodylic acid buffer, pH 6.5, in a 90% H_2_O:10% D_2_O ratio.

### UV Thermal Denaturation

UV thermal denaturation data curves were collected using a Varian Cary E3 UV-visible spectrophotometer attached to a Peltier temperature control. Each RNA sample was 10 μM in 10 mM cacodylic acid, pH 6.5, and had a layer of mineral oil on top to prevent evaporation. The samples were heated at a rate of 0.2°C/min from 25°C to 95°C with a 3 min hold. The absorbance was recorded at 260 nm, a wavelength sensitive to hairpin dissociation. (Davis et al. 1998; Draper and Gluick 1995) The data was normalized, and the first derivative processed with exponential smoothing was used to determine the melting temperature, T_m_, of each sample.

## Acknowledgements

This research was supported by NSF CHE-2029124 RAPID (MRM, JDE), NSF MRI Supercomputer CHE-1726824, NSF CHE-1950585 REU (MAS, PEL, JDE), and NIH 2R15GM127307-05 (MRM) grants.

